# Cerebral coherence in task-based fMRI hyperscanning: A new trick to an old dog

**DOI:** 10.1101/2021.07.21.452832

**Authors:** Le-Si Wang, Jen-Tang Cheng, I-Jeng Hsu, Shyhnan Liu, Chun-Chia Kung, Der-Yow Chen, Ming-Hung Weng

## Abstract

This study features an fMRI hyperscanning experiment, mapping the brains of the dyads from two fMRI sites, 305 km apart. There are two conditions: in half of the trials (the cooperation condition), the dyad had to collaborate to win and then split the reward, whereas in the other half (the competition condition), the winner took all the reward, thereby resulting in dynamic strategic interactions. Each subject took alternating turns as senders and receivers. To calculate the cerebral coherence in such jittered event-related fMRI tasks, we first estimated the feedback-related BOLD responses of each trial, using 8 finite impulse response functions (16 seconds), and then concatenated the beta volume series. With the right temporal-parietal junction (rTPJ) as the seed, the interpersonal connected brain areas in the cooperation and competition conditions were separately identified: the former condition with the right superior temporal gyrus (rSTG) and the latter with the left precuneus (lPrecuneus) (as well as some other regions of interest), both peaking at the designated frequency bin (1/16 s = 0.0625 Hz), but not in permuted pairs. In addition, the extended coherence analyses on shorter (12 s, or .083 Hz) and longer (20 s, or .05 Hz) concatenated volumes verified that only approximately in the trial length were the rTPJ-rSTG and rTPJ-lPrecuneus couplings found. In sum, our approach both showcases a flexible analysis method that widens the applicability of interpersonal coherence in the rapid event-related fMRI hyperscanning, and reveals a context-based interpersonal coupling between pairs in cooperation vs. competition.

**Author summary:** Social neuroscience is gaining momentum, while coherence analysis as one of the interpersonal connectivity measures is rarely applied to the rapid event-related fMRI. The reason could be that the inherent task design (such as the periodicity constraint for Fourier transformation), among others, limits its applicability and usage. In this fMRI hyperscanning study of a two-person strategic interactions, we independently estimated the feedback-related BOLD responses of each trial, and concatenated the beta time series for interpersonal coherence. The main advantage of this method is in its flexibility against the constraints of jittered experimental trials intermixing several task conditions in most task-based fMRI runs. In addition, our coherence results, which highlight two inter-brain couplings (e.g., rTPJ-rSTG between collaborating, and rTPJ-lPrecuneus for competing dyads) among other brain regions, plus its temporal specificity of such seed-brain couplings only between pairs, both replicate previous run-wide fMRI coherence results, and hold great promise in extending its applicability in task-based fMRI hyperscanning.

## Introduction

In recent years, social neuroscience has been one of the fast growing fields among branches of functional neuroscience, partly due to the emerging consensus about its importance in the ever changing world, from global to interpersonal context, prompting for the underlying mechanisms and ensuing remedies/interventions from the neuroscientific viewpoints. Not surprisingly, most of the queries of social neuroscience have to be ‘social’, or interactions between two or among more people or agents in nature, to be categorized as ‘social neuroscientific’. Various research paradigms, tools, and analytic methods have helped expand the breadth and depth of its inquiries. Two recent reviews summarized the growth, outstanding works, and the challenge ahead for social neuroscientists using hyperscanning [1, 2], a popular method characterized by having one or more neuroimaging devices synchronously or asynchronously, designed to make optimized social inquiries social. After listing the advent of the first hyperscanning paper [3] and ensuring achievements using various methods, one of the hidden shortcomings both reviews commonly revealed was the lack of true hyperscanning fMRI studies (and the scarce of available datasets thereof). The reason may be obvious: the technical feasibility and the collaborations among highly skilled teams required for hyperscanning were not easy for any individual fMRI labs. To fill in the gaps and limited publications in the past twenty years, we (several fMRI practitioners in southern Taiwan) have developed an internet-based hyperscanning, similar to the earlier pioneering work [4] that featured the two (or more) computers connected to a central server, receiving from and sending to client computers the signals for achieving synchronous onset triggers. By over a hundred times of running hyperscanning experiments, this fMRI study marks the fruition of years of our accumulated work.

As one of the important methods that characterizes social neuroscientific studies using hyperscanning [2, 5], cerebral coherence is a spectral measure that estimates the linear time-invariant relationship between (intra- or) inter-personal time series to generate maps of task-or condition-specific connectivity associated with seed (or electrode) regions of interest. Because of its insensitivity to slight temporal lags, which happens regularly in common social interactions, cerebral coherence has been one of the desirable choices to reveal inter-brain communications among seas of two (or more) EEG/MEG/fNIRs/fMRI signals [6–9]. However, due to both the technical challenges in conducting fMRI hyperscanning, plus the methodological constraints to meet coherence analyses (detailed next), there are so far only one [10] publication of fMRI hyperscanning studies employing coherence analysis, with a decent numbers of hyper-fNIRs, -ERP, or even -MEG studies [11–14] reporting coherence analyses. The reasons are probably due to the following: (a) cerebral coherence relies on time-to-frequency-domain (e.g., Fourier) transformation, which requires the periodicity (or fixed interval) of events/conditions to begin with. In contrast, most task-fMRI studies, especially event-related ones, are realized by irregular inter-trial intervals (ITI); (b) coherence typically requires the long task condition/runs, whereas most fMRI studies intermix several conditions of interest, each with variable numbers of blocks or short-duration (~< 20 s) trials presented in an unpredictable manner, adding another layer of complexity; (c) most importantly, despite its early onset [3], hyperscanning remains technically formidable. Some labs may have resolved the difficulty of long-distance (internet) communications by close-by fMRIs in research institutes [15] or hospitals, but the cost and benefit analysis still deter researchers from striding into these challenges.

To fill this lack of hyperscanning fMRI studies with coherence analyses, driven by the conflicts between different analysis methods and the corresponding experimental designs, we tried to make ‘having the cake and still eating it’ possible, by designing a jittered event-related hyperscanning fMRI study, adopting both the conventional GLMs, other derived analyses such as connectivity and multivariate mappings [16], and cerebral coherence analyses in the present study. To do so, Turner et al [17] that also incorporates the iterative least squares-separate method (the Turner method hereafter, similar to that of Mumford et al. [18]), convolved with the stick-like finite impulse series to estimate feedback-related time courses data of various TRs (e.g., 6, 8, and 10). What distinguishes the two methods was that the method of Mumford et al. was to convolve with a hemodynamic response function (HRF) function, whereas the Turner method was to convolve with one stick function per TR.

Therefore, by adopting the Turner method that transformed the original event-related fMRI dataset into the TR-beta-concatenated time series of data points ready for coherence analysis. To test and validate this transformed data series, a task-based fMRI experiment was implemented while subjects cooperated (and split the reward) or competed (and the winner took all the reward) in a strategic cheap game [19]. The feedback period of the cooperation condition was especially of interest, since it is the time during which the dyads verified and strengthened their mutual understanding. Significant increases in interpersonal coherences between the rTPJ and the rSTG were identified, resembling past findings reported in the relevant literature [10, 15, 20, 21]. That is, there was significant interpersonal coherence between the rTPJ and the rSTG in the cooperation feedbacks, found only within communicating pairs (but not among random pairs). To our knowledge, this is the first study running coherence analysis on a task-based jittered event-related fMRI experiment. It therefore expands the range which the coherence analysis could be applied to while extra information is added alongside or even the original GLM or other types of task-based fMRI analysis methods are combined with.

## Results

The experimental task of this hyperscanning fMRI study was a strategic cheap talk game with two conditions: cooperation (winners divided the $200 reward) vs. competition (the winner took all the $150 reward). Both players took turns as senders and receivers alternatively (see Figs 1A-C for details of the experimental design).

**Fig 1.**
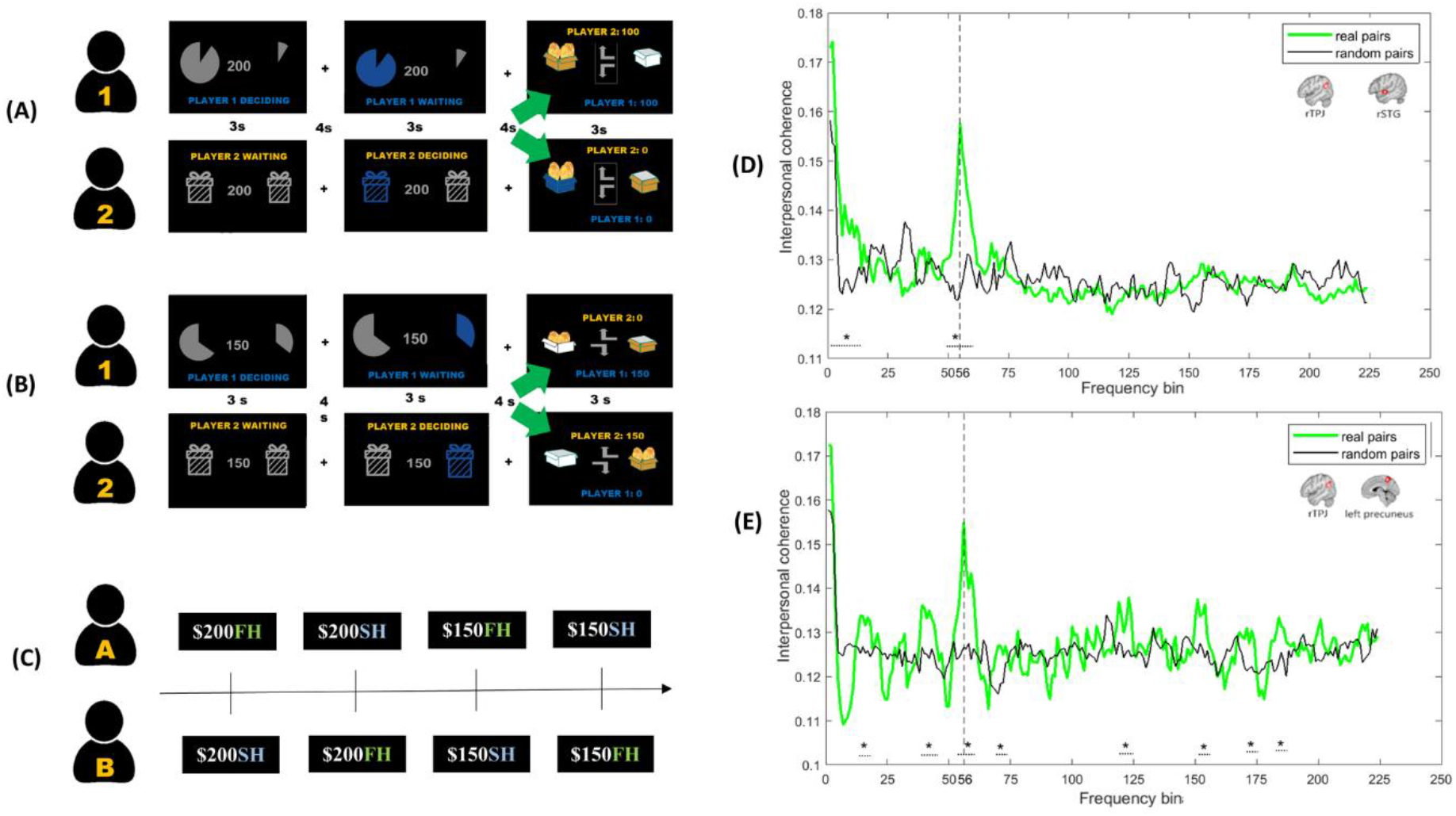
The present hyperscanning fMRI experiment of the strategic game with two conditions. Player 1 and Player 2 alternatively served as the sender and the receiver. In both conditions, the sender, knowing the probability of the treasure box, first suggested a box for the receiver to choose. Then, the receiver made the decision of which box was opened. (A) In the cooperation condition (NT$ 200), the dyad split the reward when the box with money was chosen correctly. (B) In the competition condition (NT$ 150), either the sender or the receiver won the reward when the result was revealed. In the present study, we only analyzed the interpersonal coherence during feedback time of the cooperation condition, which was the event when the result of the correct box was revealed (15-17 s). (C) A 2-by-2 manipulations of “cooperation (or NT$ 200) vs. competition (or NT$ 150)” and “sender (first-hand, or FH) vs. receiver (second-hand, or SH)” in the alternating fashion, arranged in a $200FH-$200SH-$150FH-$150SH order for Player A and in a $200SH-$200FH-$150SH-$150FH order for Player B. (D-E) Interpersonal coherence spectra between the rTPJ and the rSTG (D) and between the rTPJ and the left precuneus (E) of real communicative pairs (shown in green) as well as random pairs (shown in black) (\**p* > 0.01 and three consecutive bins). The dominant (or peak) frequency, 0.0625 Hz (1/16 s), located at the 56^th^ frequency bin (along the x axis), revealed the 2^nd^ highest coherence (the first bins being around time zero). This bin corresponded to the average trial frequency (once every 17 s), as well as to the concatenated event frequency (beta series built by combining 8 betas, every 2 s/ 1 TR, after the trial feedback time); therefore, the heightened coherence between rTPJ-rSTG (D) and rTPJ-lPrecuneus (E) suggest that the dyads reached certain degrees of synchronization.

### Interpersonal and inter-regional coherence in the cooperation and the competition condition

Figs 1D and 1E represent the average cerebral coherence between 33 pairs of the interpersonal rTPJ and rSTG (Fig 1D, cooperation condition), and rTPJ and lPrecuneus (Fig 1E, competition condition), based on the feedback-initiated time series (8 TR/volumes) modeled by the finite impulse response (FIR) deconvolutional analysis, or the Turner method. Especially noteworthy is the resemblance of the current Figs 1D and 1E to the similar figure of an earlier study (cf. with Fig 2C in Stalk et al., 2014 PNAS, pp. 18185 [10]). This coherence peak was located at the 56^th^ frequency bin (out of 448/2 = 224 bins), corresponding to 0.0625 Hz (1/16 s), and exactly matched the concatenated trial periodicity. These coherence peaks were identified only within the communicating (true) pairs, but not in the permuted pairs (e.g., black lines in both Figs 1D and 1E). This is surprising given that the real and permuted pairs were identical in many aspects: the trial orders in each run, the event frequency across all 4 runs, and subjects’ response time ranges, etc., except the only difference being the shared interaction history of the real pairs. In other words, only in the simultaneously cooperating/competing pairs did the interpersonal rTPJ-rSTG/-lPrecuneus coherence emerge, but not in randomly matched pairs where each was drawn from a different session. In addition, our finding also echoes the importance of STG in social interaction. Only those with shared communication history, i.e., truly interacting pairs, were seen to have more coherent activities in the STG. Moreover, the comparison between findings in cooperation and competition conditions signifies that the STG plays an important role in social interaction only when the communication results were mutually accepted, i.e., in cooperation, but not when in competition.

**Fig 2.**
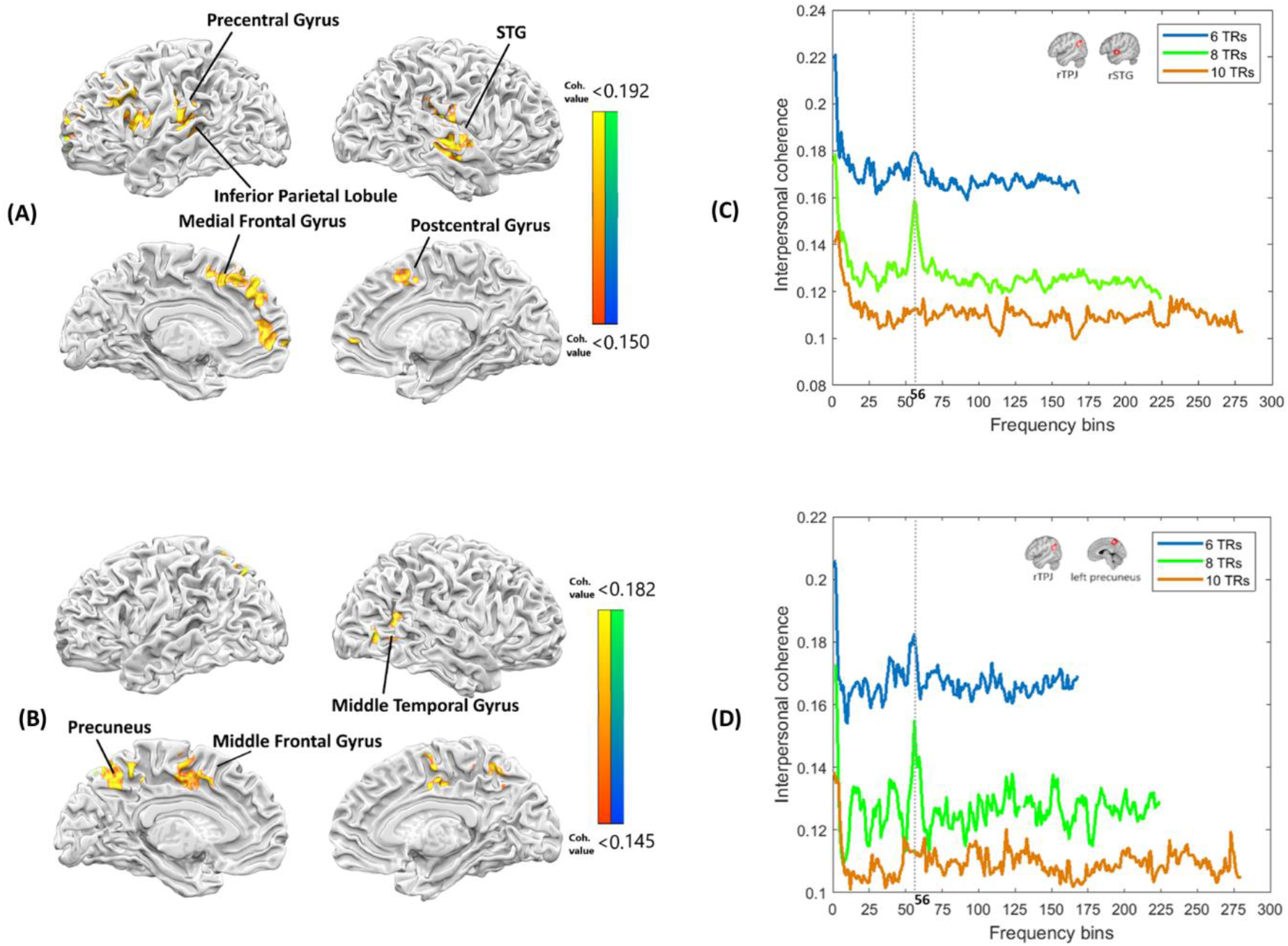
rTPJ-to-whole-brain maps and seed-to-whole-brain coherence mapping with 3 different TRs of beta volumes in cooperation and competition. (Figs 2A and 2B) Brain regions showing significant coherence with the rTPJ during the event of feedback time. (A) In the cooperation condition, under the cluster k-threshold of 40 voxels and the applied map threshold of 0.15, the ROIs with significant coherence with the rTPJ in the cooperation condition were: the postcentral gyrus, the rSTG, the left medial frontal gyrus, the left precentral gyrus, the left superior frontal gyrus, and the left inferior parietal lobule. (B) In the competition condition, under the cluster k-threshold of 40 voxels and the applied map threshold of 0.145, the coherent ROIs were the left precuneus, the right middle temporal gyrus and the left middle frontal gyrus. (Figs 2C and 2D) Seed-to-whole-brain coherence mapping with the rTPJ with 6, 8, and 10 beta volumes. (C) In the cooperation condition, the interpersonal coherence between the rTPJ and rSTG was presented. The target frequency bin (1/16 s = 0.0625 Hz, found at the 56^th^ frequency bin) of 8 beta volumes significantly peaked, while that of 6 or 10 (both found at the 56^th^, too) was relatively noisier. (D) In the cooperation condition, the interpersonal coherence between the rTPJ and the left precuneus was presented. The left precuneus at the target frequency bin of 8 beta volumes (1/16 s = 0.0625 Hz, found at the 56^th^ frequency bin) was significantly coherent with the rTPJ. The interpersonal coherence of the 56^th^ with 6 or 10 beta volumes, though significantly different against the values from the permuted pairs, was nonetheless noisier relative to the 8 TRs.

### Seed-to-whole brain coherence

Figs 2A and 2B show the seed (rTPJ) to the whole-brain interpersonal coherence map, averaged across 66 participants, on cooperation and competition conditions, respectively. To be clear, these seed-to-brain coherence analyses were done prior to the seed-to-ROI coherences shown in Figs 1D and 1E (though reversed in the presentation order). As shown in Fig 2A, medial frontal gyrus, inferior parietal lobule, postcentral gyrus, and precentral gyrus, other than the rSTG (Fig 1D), were among the brain regions that showed high interpersonal coherence with the seed rTPJ. Likewise, in Fig 2B, middle temporal and middle frontal gyrus, other than the precuneus (Fig 1E), were among the regions showing higher interpersonal coherence with the seed rTPJ. S1 Table provides more details of these clusters. These regions may be among the networks subserving the dyad interactions for both cooperation and competition. To be comprehensive, the complete whole-brain coherence maps of 224 frequency bins were integrated into two separate movies (see S4 Fig, for cooperation and competition conditions, respectively).

### Comparing coherences across different trial durations

To clarify whether the seed-to-brain coherence (Figs 2A and 2B) results were caused more by the 8-TR (16 s) concatenation into the event time series, or more by the close match to the average trial frequency (17 s per trial), here we created another two concatenated volume series with 6 and 10 TRs, or 12 and 20 seconds each, and repeated the coherence analyses. Our rationale was that, if the 8-TR (16 s) coherence result was due more to the concatenated volume periodicity, then one should observe the same inter-regional coherences in the same 6-TR (or 1/12 s, 0.083 Hz) and 10-TR (1/20 s, 0.05 Hz) cases. In contrast, if the 8-TR coherence results were due more to the average trial frequency in the real fMRI data, then 6-TR or 10-TR cases should not yield the desired, or less strong, coherence, especially around the target (e.g., 56^th^) frequency bin. The additional rTPJ-ROI coherence analyses were carried out and plotted above/below the original 8-TR (16 s) coherence plots (Figs 2C and 2D, for cooperation and competition conditions, respectively) for comparison purposes. The results clearly favor the contribution by the average trial frequency of the real data, rather than that by the right number of concatenated volumes, in helping shape the current coherence results. These analyses not only lend further support to our methodology, but provide additional consideration in the coherence analysis protocol (e.g., to match the average trial frequency).

## Discussion

This study aims to uncover inter-brain coherences using fMRI hyperscanning, a method that has shown great promise in helping shape the inquiries and paradigms of social neuroscience. Despite its technical challenges, recent breakthroughs [2] have paved ways to realize complexity underlying social dynamics, a critical component toward better understanding of human lives, as well as interactions with the earth, and the future prospect of both. As one conspicuous limitation to the application of interpersonal coherence in fMRI hyperscanning, in order to fit the periodicity requirements of frequency decompositions, the experimental design has to be adapted into regular, periodical, and full with interactions [10, 22]. Such limitations not only reduce the range of available experimental paradigms, but also leave the interpretations to coherence-only dimensionality. To fill in the gap and extend the data analysts’ repertoire, here we report a jittered-erfMRI hyperscanning experiment, in which the trial-feedback responses were TR-based deconvolved into 8-volume series. With the concatenation of cooperation and competition conditions separately across runs for each participant, the final 448 (8 volumes x 14 trials x 4 runs) time series were ripe for pair-specifically or pair-randomly coherence analyses. Paralleling the past studies [10, 22], the coherence analyses on these transformed dataset yield comparable results with Fig 2C in Stalk et al. [10]. In a way, the realization of ‘having both cakes and eating it’ is compellingly demonstrated, if not mentioning the additional task-based fMRI analysis, which was reported elsewhere [16].

The reported inter-personal rSTG-rSTG coherence is what Stalk et al. in 2014 [10] attributed as the source of emergent ‘meanings’. Our findings of both the rSTG-rSTG (see S7 Fig), as well as the rTPJ-rSTG (see Fig 1C) coherence in the cooperation condition, along with rTPJ-lPrecuneus, rSTG-other target regions in the competition conditions, both point to the general patterns of reward, execution, and theory-of-mind (ToM), -related networks. The presence of converging rSTG-rSTG findings is encouraging, providing extra support for the notion of rSTG as the target of ‘conceptual alignment’ [22], but the implication behind rTPJ-rSTG is less clear. Here we propose that even though the rTPJ, the alluded target area for ToM, would be less of a degree to the level of ‘mutual understanding,’ the rTPJ-rSTG coherence still represents a second best approximation of mutual rapport. Under the 7 computations behind social neuroscience [23], this could be akin to the social perception and social inference stages that underlie social trust/rapport.

Prior studies regarding interpersonal interactions have adopted various methods investigating the role of rTPJ and related brain areas (such as rTPJ/STG/superior temporal sulcus, STS). For example, Abe et al. [24] adopted multivariate autocorrelation models to assess root mean square error (RMSE) between hyperscanning dyads’ grip forces in collaboration. Bilek et al. [15] used groupwise Independent Component Analysis (ICA) for identifying the rTPJ as sources of important components for collaborating fMRI pairs during hyperscanning, and the same fMRI data were later reanalyzed with pairwise directional coherence [20], and found neural coupling through the rTPJ to the ventromedial prefrontal cortex (vmPFC) for feed-forward, and the posterior cingulate cortex (PCC) to the right STS for feed-backward, information exchanges. Using fNIRS, Tang et al. [25] found increased interpersonal brain coherences during face-to-face economic exchange between pairwise rTPJs, highlighting its importance in collaborative social interactions. With EEG-hyperscanning, Jahng et al. [21] provided evidence in neural dynamics between dyads when nonverbal cues involving the rTPJ were adopted to predict opponents’ intentions to cooperate or defect during the face-to-face prisoner’s dilemma game. All the above-mentioned studies adopted either specialized analysis methods or additional cues, such as eye contacts [15] and facial co-analysis between patients and clinicians [26] in their analyses of the information flow. The common mechanism of interpersonal rapport triggering similar brain activities should be a common theme [27–29]. All in all, with the recent literature that suggests the importance of neuron oscillations at specific frequencies [30, 31] or decreased gamma-band in the competitive pairs [32], all lend support to the ‘communication-through-coherence’ hypothesis [33]. Therefore, coherence between two neuronal groups, brain areas, pairs, organizations, societies, or even countries or cultures could be exploited to reach effective communication, benefiting the future of human societies.

One recent comment with the title ‘Hyperscanning: beyond the hype’ of coherences in hyperscanning fEEG/fNIRS studies’, listed several cautionary points that challenged the interpretations of cross-brain coherence data [34]. Indeed, coherence may still be epiphenomenal given that fMRI has been traditionally considered to be a ‘correlational research tool’; meanwhile hypotheses would be best corroborated by the Multi-Brain Stimulation (or MBS) methods [35]. That is, as the current study exemplifies on the availability of coherence with the typical task-based fMRI, hyperscanning fMRI, with its technical advancements and methodological improvement [2], will possibly even correlate with subject/condition-wise variables, continue to thrive for significant breakthroughs, and uncover more neural underpinnings of common or subtle within- or between-individual/group dynamics.

## Materials and Methods

### Participants

Thirty-three pairs of participants between 20 and 30 years of age (M = 23.4, SD = 2.9) were recruited from National Taiwan University (NTU) and National Cheng Kung University (NCKU), situated in northern and southern Taiwan, respectively (305 km apart). All participants are native Taiwanese speakers, with normal or corrected-to-normal vision, and reported no history of psychiatric or neurological disorders. Participants gave written informed consent, and adhered to the relevant guidelines and regulations approved by the NCKU Governance Framework for Human Research Ethics https://rec.chass.ncku.edu.tw/en, with the case number 106-254.

The experimental task was a strategic cheap talk game with two conditions: cooperation vs. competition. Participants were assigned alternatively as senders or receivers. The goal of the participants was to choose the treasure box with money to get the reward. Both of the conditions started with the sender first suggesting a box and then the receiver deciding. In the cooperation condition, the dyad then split the $200-reward if the receiver opened the correct box. In the competition condition, only the receiver (sender) got the $150 reward if the receiver opened the correct (incorrect) box. Each trial took 17 seconds to complete: 3 seconds for the sender’s decision, 4 seconds of fixation, 3 seconds for the receiver’s decision, 4 seconds of fixation, and 3 seconds of feedback respectively. The inter-trial interval was approximately 3 to 9 seconds (see Figs 1A and 1B). The period between 15 to 17 seconds (feedback time, revealing the result of the correct treasure box) was the observation vector for our coherence analysis, since it was the time the dyad built up the cooperative/competitive mindset regarding their opponent. We extracted 8 beta series from feedback time to be classified by using a finite impulse response function to extract jittered event-related activity estimates for each of 4 runs in this strategic game and concatenated the beta series in a periodic data format. We then conducted a GLM-based multi-parameter method iteratively to allow trail-by-trail estimation. Finally, we conducted a time-frequency analysis to present the interpersonal coherence between subjects. In order to run seed-brain coherence analysis of Fieldtrip in jittered, rapid-event experiments, the preprocessing job for the dataset was demonstrated as follows.

### Experimental task

In the present hyperscanning fMRI experiment, also known as the ‘Opening Treasure Chest game’. a 2-by-2 manipulations of “cooperation (or NT $200) vs. competition (or NT $150)” and “sender (first-hand, or FH) vs. receiver (second-hand, or SH)” were administered in the alternating fashion (Fig 1C). To balance the expected utility of both conditions, the cooperation condition (NT $200) was set to have a 75% success rate, as the players were expected to split the $150 reward in the competition condition. Therefore, in the ideal situation where the dyads were 100% trusting in the cooperation condition, the expected utility for each person is ($200*75%) /2 = $75, whereas in the competition condition, the expected utility for each is also the same ($150*50% = $75). There are 28 trial events in each run, and the conditions ($200 vs. $150) alternated between pairs of trials, and subjects changed their roles after each trial (arranged in a $200FH-$200SH-$150FH-$150SH order for Player A and in a $200SH-$200FH-$150SH-$150FH order for Player B, repeated for 7 times), and four runs for each pair. The subject’s role on either site as player A or B (Fig 1C) was pseudo-randomly assigned while equally presence on both roles on both sites was ensured. As depicted in Figs 1A and 1B, at the beginning of each trial there were two boxes presented, only one containing the target reward. The objective was to choose the correct box so to either split ($200) or take all ($150) the reward. As the sender, the participant alone was informed of the probabilities of money in both boxes and was told to suggest which box the receiver should choose. After viewing the suggestion from the sender, the receiver selected which box to open. In all trials, the box chosen by the receiver is the final choice of both the sender and the receiver. In the cooperation condition, the dyad splits the $200-reward if the chosen box was with money. In the competition condition, the receiver got what was in the chosen box while the sender got the other one. Only one of them got the $150 reward. The participants were offered a financial payment of NT $600 (~US $20), plus a bonus depending on the rewards they have won from one randomly selected trial.

### fMRI data acquisition and preprocessing

The fMRI images of senders and receivers were acquired simultaneously in two MRI scanners, one in NCKU Tainan, another in NTU Taipei, 305 km apart. The MRI scanner at NCKU Mind Research and Imaging Center is a 3-Tesla General Electric Discovery MR750 (GE Medical Systems, Waukesha, WI), equipped with an 8-ch head coil. Whole-brain functional scans were acquired with a T2* EPI (TR = 2 s, TE = 33 ms, flip angle = 90°, 40 axial slices, voxel size = 3.5 × 3.5 × 3 mm^3^). High-resolution T1-weighted structural scans were acquired using a 3D fast spoiled grass (FSPGR) sequence (TR = 7.65 ms, TE = 2.93 ms, inversion time = 450 ms, FA = 12°, 166 sagittal slices, voxel size = 0.875 × 0.875 × 1 mm^3^). Another fMRI scanner, which is located at NTU (Imaging Center for the Body, Mind, and Culture Research) is a 3-Tesla PRISMA (Siemens, Erlangen, Germany) scanner equipped with a 20-channel phase array coil. Whole-brain functional scans were acquired with a T2*-weighted EPI (TR = 2 s, TE = 24 ms, flip angle = 87°, 36 axial slices, voxel size = 3 × 3 × 3 mm^3^). High-resolution T1-weighted structural scans were acquired using a MP-RAGE (TR = 2.0 s, TE = 2.3 ms, inversion time = 900 ms, FA = 8°, 192 sagittal slices with 0.938 × 0.938 × 0.94 mm^3^ voxels without an interslice gap). The fMRI data were preprocessed and analyzed using BrainVoyagerQX v. 2.6 (Brain Innovation, Maastricht, The Netherlands) and NeuroElf v1.1 (https://neuroelf.net). After slice timing correction, functional images were corrected for head movements using the six-parameter rigid transformations, aligning all functional volumes to the first volume of the first run. High-pass temporal filtering (with the default BVQX option of GLM-Fourier basis set at 2 cycles per deg, but no spatial smoothing) was applied. The resulting functional data were co-registered to the anatomical scan via initial alignment (IA) and final alignment (FA), and then both functional (*.fmr) and anatomical (*.vmr) files were transformed into Talairach space.

### Data Preparation and the coherence analysis

The procedure of converting an event-related jittered fMRI dataset into a format ready for coherence analysis is detailed in Fig 3. First, the feedback period of any given trial (here the cooperation, or $200, condition, was used for illustration (Fig 3A). Next, as shown in Fig 3B, each run was decomposed into con(dition)_200_1st(hand), con_200_2nd(hand), con_150_1st, con_150_2nd, others (missing trials due to subject oversight, internet problem, etc.), and more importantly here, the two feedback conditions, fb_200 (the cooperation condition) and fb_150 (the competition condition). There were 4 runs, with 28 trials each (14 for the cooperation, and 14 for the competition condition).

**Fig 3.**
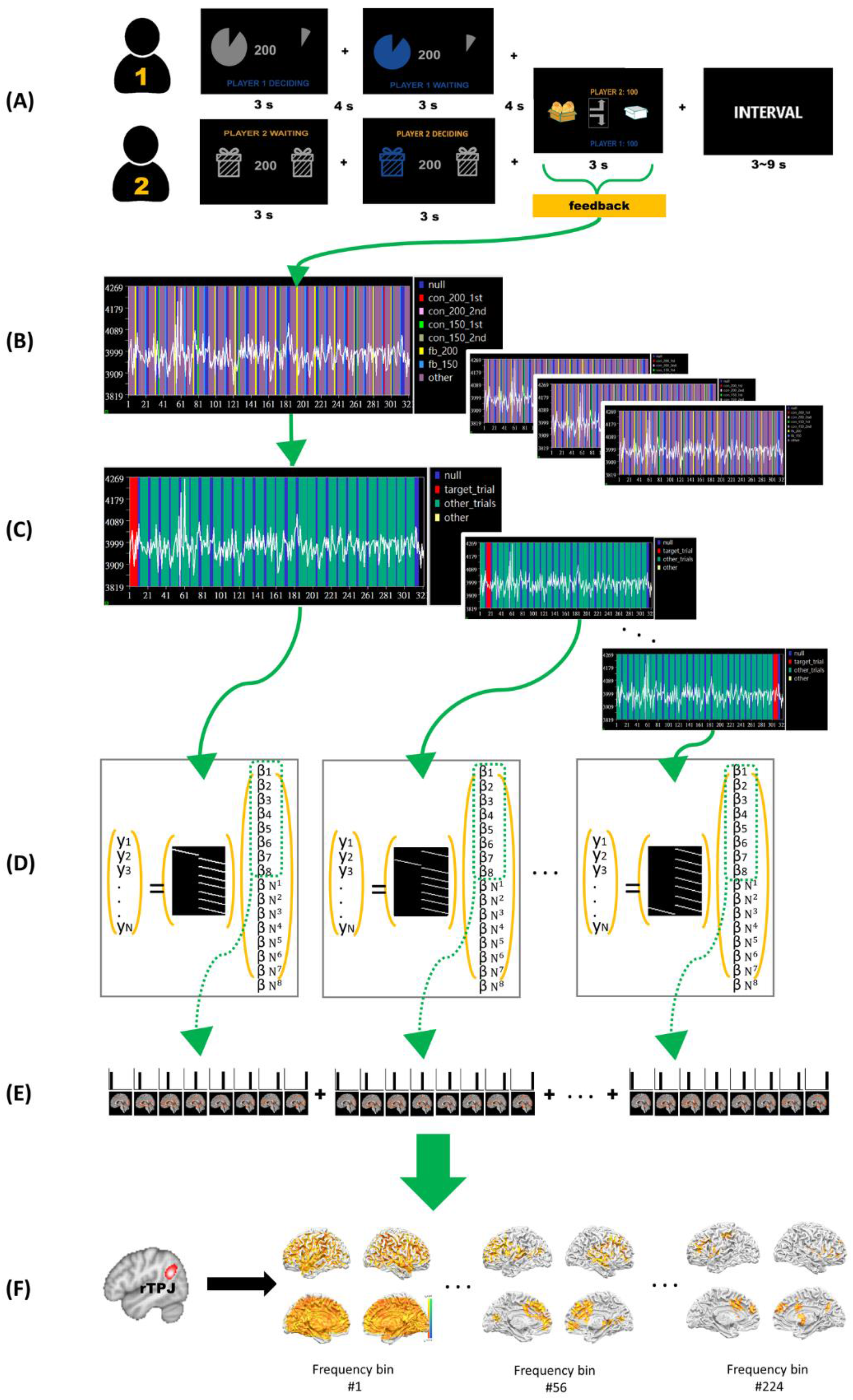
The schematic procedure of converting an event-related jittered fMRI dataset into a format ready for coherence analysis. (A) “Feedback”: defined as the time between the onset when the result of the box-opening game was revealed, to the end of the 3-s period, applied to both the cooperation ($200) and the competition ($150) trials (14 in each run). (B) The stimulus design matrix (or SDM file) of a run (totally 4 runs), along with the voxel time series, was presented: each run contained 7 regressors. (C) Converting a 7-regressor protocol response time (or PRT file) into a two-regressor version, with “target_trial” and “other_trials” looping iteratively across all the chosen trials. For example, when the 1^st^ trial was the target, the other trials included 2^nd^ trial to the last trial; when the 2^nd^ trial was the target, the other trials were the 1^st^, and the 3^rd^ to the end; and so on; (D) The design matrix for a single deconvolutional GLM was created, with 8 stick-like extensions for each of the two regressors (“target_trial” and “other_trials”). This method allowed a least-square separation of the target trial, from which the whole brain beta series were independently estimated subsequently for the 8 TRs following the feedback-initiated event (cooperation condition here); (E) Concatenating all the 8 TRs from each trial, resulting in 448 volumes (8 TRs x 14 trials x 4 runs), as the periodic data format ready for coherence analysis; (F) The rTPJ was the seed ROI for the interpersonal seed-brain coherence analysis; thus the regions with high coherence were mapped.

The protocol (or .prt file) of a fMRI run (totally 4 runs), along with the voxel time series, was created: each run contained 7 regressors, con_200_lst, con_200_2nd, con_150_1st, con_150_2nd, fb_200, fb_150, and others (Fig 3B). The third step, which was the core of Turner method, was to convert a 7-regressor prt into a two-regressor version, with “target_trial” and “other_trials” looping iteratively across all the chosen trials (Fig 3C). For example, when the 1^st^ trial was the target trial, the other trials included 2^nd^ trial to the last trial; when the 2^nd^ trial was the target trial, the other trials were the 1^st^, and the 3^rd^ to the end, and so on. With the rearrangement of the regressors, we created the design matrix for a single deconvolutional GLM, with 8 stick-like extensions for each of the two regressors (“target_trial” and “other_trials”) (Fig 3D) [17]. This method allowed a least-square separation of the target trial, from which the whole brain beta series were independently estimated subsequently for the 8 TRs following the feedback-initiated event. Concatenating all the 8 TRs from each trial resulted in 448 volumes (8 TRs x 14 trials x 4 runs) (Fig 3E), as the periodic data format ready for coherence analysis. To mend the missing trials, we then manually calculated approximate time and replaced the missing time slots for the later need of coherence data structure, instead of padding zeros, which greatly and statically affects the frequency domain. The rTPJ (and rSTG) was chosen as the seed ROI for the interpersonal seed-brain coherence analysis; thus the regions with high coherence were separately mapped (in Figs 1D and 1E and S7 Fig, respectively). In addition to the ROI-ROI coherence value, another index of Fourier decomposition, the phase-locked value (PLV), was also mapped (see S3 Fig). As seen, the PLV also supports the notion of interpersonal coherence. Coherence analysis was performed using the FieldTrip toolbox, a MATLAB software toolbox for MEG, EEG and iEEG analysis, such as time-frequency analysis (https://www.fieldtriptoolbox.org/). Whole-experiment BOLD signal time series were extracted from regions of interest, synchronized to experiment cooperation feedback time as the observation vectors. We extracted the beta series from the desired events built by every 2 seconds (1 TR), with 8 betas for each event, and concatenated into 8 x trial numbers (4 runs and 14 trials each run here) of data for coherence analysis. For each observation vector, the time series was collapsed into multiple consecutive overlapping windows of 448 volumes (75% overlap). The windows were then tapered with a set of Slepian tapers before spectral estimation and calculation of the magnitude squared coherence. This resulted in a spectral smoothing of 0.005 Hz.

### Interpersonal seed-brain coherence analysis

We revised the codes from the Fieldtrip website (https://www.fieldtriptoolbox.org/example/correlation_analysis_in_fmri_data/). First, the groupwise whole brain mask, derived from averaging over the whole 66 participants, were 53,207 voxels, and was divided into 54 segments (every 1,000 voxels for each, and the 54^th^ with 207 voxels.) Next, the beta values from the seed ROI (rTPJ) from the whole brain values. We averaged the rTPJ values by subjects. The input data for Fieldtrip’s frequency analysis should be organized in a structure, label, time, and trial to perform preprocessing. Afterwards, we saved their coherence values between the rTPJ and the whole brain. In the structure of Fieldtrip coherence analysis was found the frequency time series of the 224 frequency bins (out of 0.5/224, fMRI sampling frequency) / 448/2 (Nyquist equation) = 224 frequency bins). Since we had 8 beta series (each for 2 seconds), we thus chose the 56^th^ (1/16 s = 0.0625 Hz) as our designated frequency of interest.

With the 54 segments of coherence values concatenated, we created a whole-brain map for each pair separately based on the cooperation condition and the competition condition. Most importantly, we were able to designate the frequency bin we would like to explore (Fig 3F). Then, we concatenated all the whole brain maps from the same condition, and the condition-based whole-brain mapping was ready to run analyses via NeuroElf, a MATLAB toolbox supposed to facilitate a subset of neuro-functional imaging applications.

From the results of clustered maps, with the cluster k-threshold over 40 voxels and the applied map threshold of coherence value (0.15), the ROIs with significant coherence in the cooperation condition were listed; postcentral gyrus, the rSTG, the left medial frontal gyrus, the left precentral gyrus, the left superior frontal gyrus, and the left Inferior Parietal Lobule (Fig 2A and S1 Table). In the competition condition, under the same cluster k-threshold over 40 voxels and the applied map threshold of coherence value (0.145), the coherent ROIs were the left precuneus, the right middle temporal gyrus and the left medial frontal gyrus (Fig 2B and S1 Table). In this way, we were able to exemplify the maps of the ROIs we would like to investigate further. We chose the rSTG in the cooperation condition for its association with mutual understanding, and the left precuneus in the competition condition for its increases in brain activity during social competitions.

### Statistical inference at the group level

Pair specificity of BOLD signal synchronization was tested by comparing interpersonal cerebral coherence calculated on BOLD signals of participants forming real pairs (real pairs = 33) with cerebral coherence calculated on BOLD signals of participants that did not share a communicative history (e.g., The sender from the first pair with the receiver from the second pair and vice versa; Random pairs = 2112; i.e., 33 × 32 x 2 combinations). The coherence measures were entered into a second-level random-effects analysis correcting for multiple comparisons at the cluster level (\**p* < 0.01; 1,000 randomizations across participant pairs).

## Experiment design

JTC, IJH, DYC, and MHW; fMRI hyperscanning debug and data collection: JTC and DYC; Data analysis: MHW, LSW, and CCK; manuscript preparation and final approval: LSW, SL, CCK, DYC, and MHW.

## Abbreviations

BOLD: blood-oxygen-level-dependent
EEG: electroencephalography
ERP: event-related brain potential
FH: first-hand
FIR: finite impulse response
fMRI: functional magnetic resonance imaging
fNIRs: functional near-infrared spectroscopy
GLM: general linear model
HRF: hemodynamic response function
ICA: independent component analysis
ITI: irregular inter-trial intervals
lPrecuneus: left precuneus
MBS: multi-brain stimulation
MEG: magnetoencephalography
NCKU: National Cheng Kung University
NTU: National Taiwan University
PLV: phase locked value
PCC: posterior cingulate cortex
PRT: protocol response time
RMSE: root mean square error
ROI: region of interest
rSTG: right superior temporal gyrus
rTPJ: right temporal parietal junction
SDM: stimulus design matrix
SH: second-hand
STS: superior temporal sulcus
ToM: theory-of-mind
TR: time of repetition
vmPFC: ventromedial prefrontal cortex

## Acknowledgments

We thank the NCKU Mind Research and Imaging Center (MRIC) at NCKU, and the imaging center for integrated Body, Mind, and Culture research at the NTU, for the generous support and equipment availability for this collaboration. Special thanks go to Siao-Shan Shen and HanShin Jo for comments on the analysis code, and all the hyperscanning team for carrying out the experiment, as well as the preprocessing of fMRI data.

## Supporting Information

**S1 Table.** rTPJ-Brain interpersonal coherence in cooperation and competition.

| Cooperation condition with 448 beta volumes |  |  |  |  |  |
| --- | --- | --- | --- | --- | --- |
| Cluster | Anatomical region | x | y | z | coherence value |
| Cluster 1 (voxel: 59) | R Postcentral Gyrus | 56 | -20 | 17 | 0.1885 |
| Cluster 2 (voxel: 51) | R Superior Temporal Gyrus | 53 | -24 | 3 | 0.1852 |
| Cluster 3 (voxel: 65) | L Medial Frontal Gyrus | -15 | 57 | 10 | 0.181 |
| Cluster 4 (voxel: 85) | L Precentral Gyrus | -58 | 10 | 12 | 0.1751 |
| Cluster 5 (voxel: 59) | L Superior Frontal Gyrus | -8 | 39 | 42 | 0.1711 |
| Cluster 6 (voxel: 58) | L Inferior Parietal Lobule | -55 | -25 | 27 | 0.1683 |
| Competition condition with 448 beta volumes |  |  |  |  |  |
| Cluster | Anatomical region | x | y | z | coherence value |
| Cluster 1 (voxel: 66) | L Precuneus | -3 | -49 | 49 | 0.1762 |
| Cluster 2 (voxel: 46) | R Middle Temporal Gyrus | 40 | -64 | 9 | 0.1684 |
| Cluster 3 (voxel: 56) | L Middle Frontal Gyrus | -21 | -1 | 46 | 0.164 |

rTPJ-Brain interpersonal coherence in the cooperation condition [Coherence value threshold > .15, cluster size > 40 voxels threshold] and in the competition during the feedback time with 448 beta volumes [Coherence value threshold > .145, cluster size > 40 voxels threshold].

**S2 Fig.**
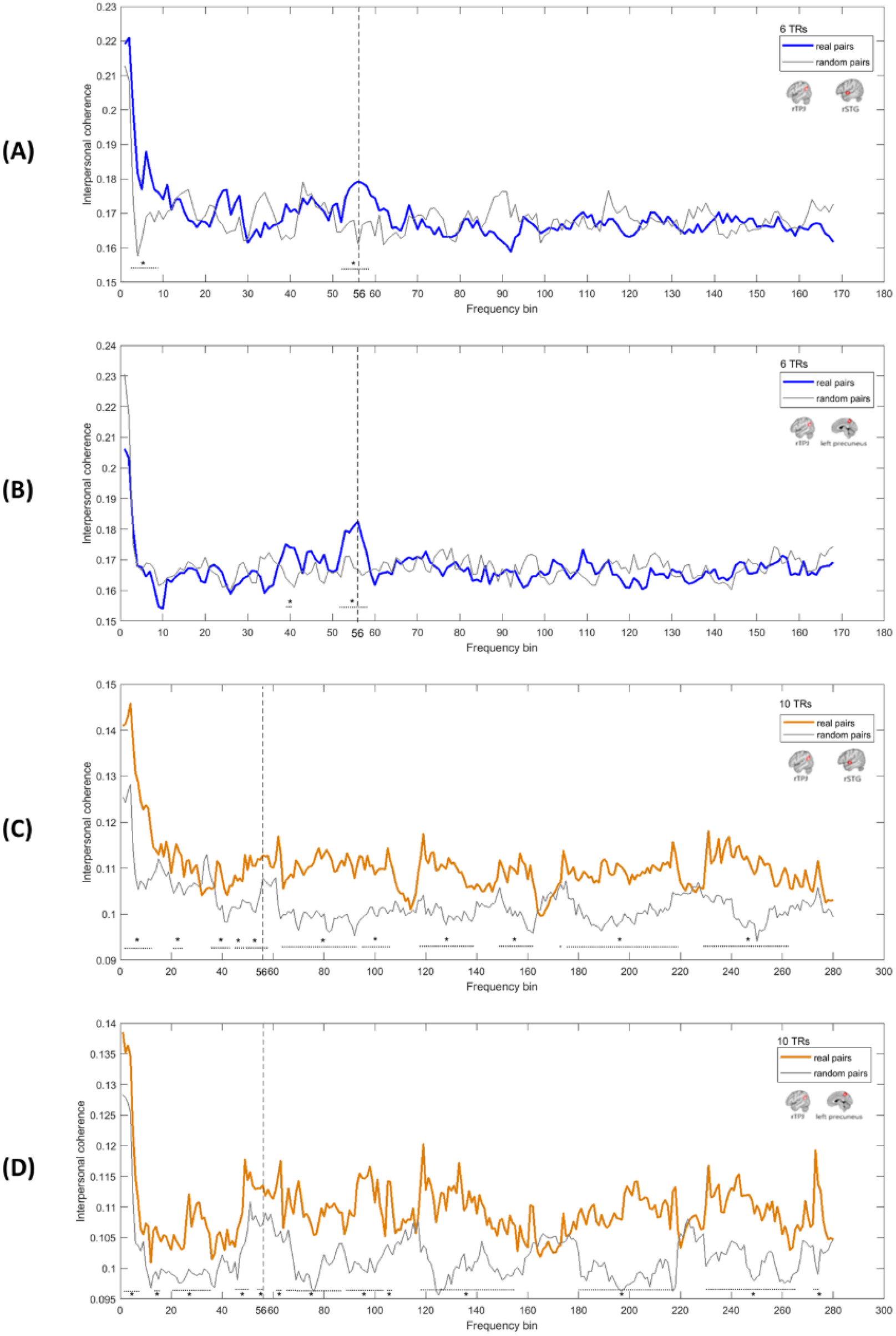
Interpersonal coherence spectra between the rTPJ and the rSTG and between the rTPJ and the left precuneus of real communicative pairs. The observed frequency was located at the 56^th^ frequency bin (along the x axis). (A and B) This bin corresponded to the average trial frequency (once every 12 s), as well as to the concatenated event frequency (beta series built by combining 6 betas, every 2 s/ 1 TR, after the trial feedback time), with coherence between rTPJ-rSTG (A) and rTPJ-lPrecuneus (B). (C and D) This bin corresponded to the average trial frequency (once every 20 s), as well as to the concatenated event frequency (beta series built by combining 10 betas, every 2 s/ 1 TR, after the trial feedback time), with coherence between rTPJ-rSTG (C) and rTPJ-lPrecuneus (D) (\**p* < 0.01 and 3 consecutive bins).

**S3 Fig.**
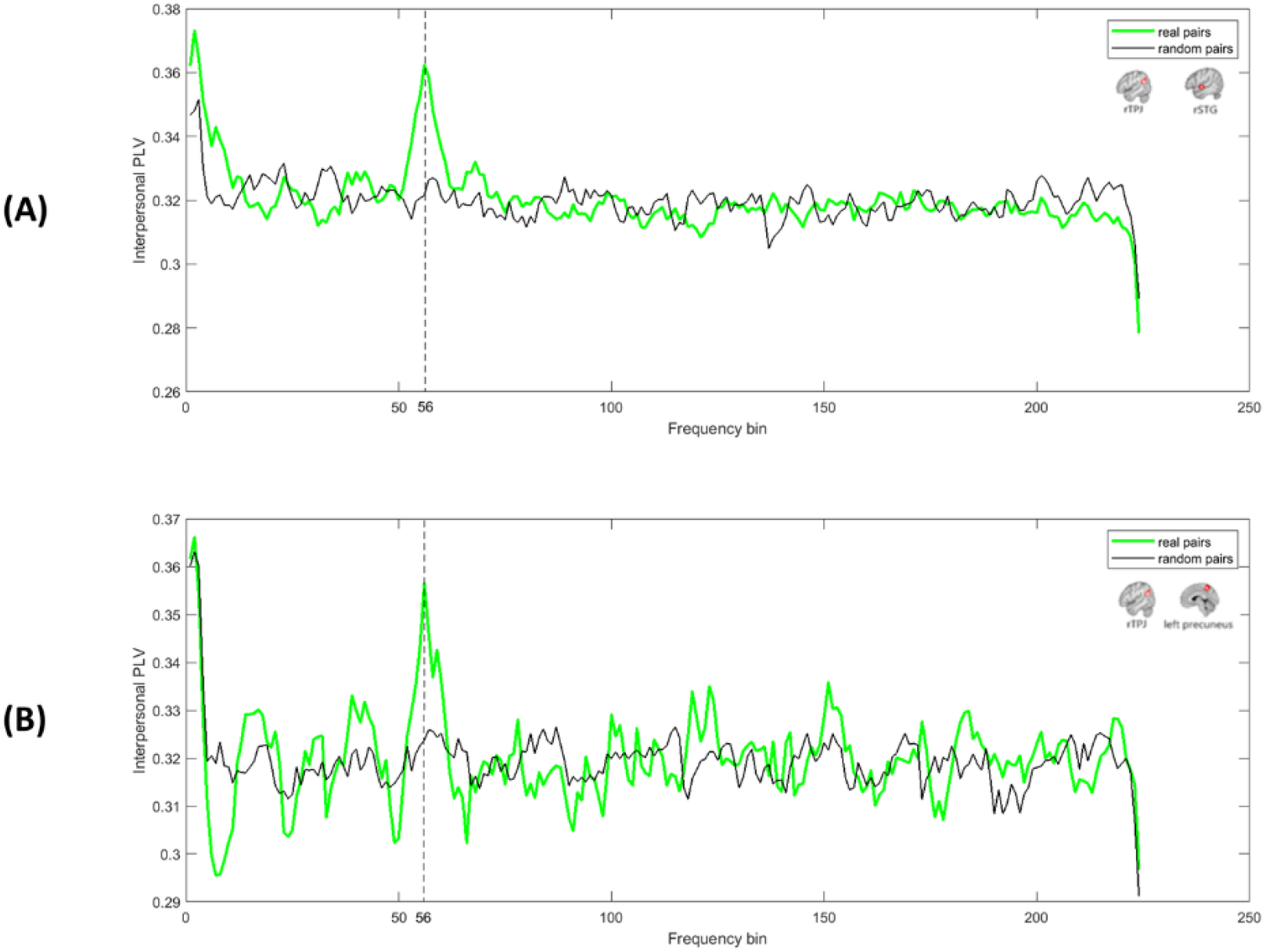
Phase-locking value with the observed frequency was located at the 56^th^ frequency bin (along the x axis). (A) PLV between rTPJ-rSTG for the cooperation condition. (B) PLV between rTPJ-lPrecuneus for the competition condition.

**S4 Vedio.**
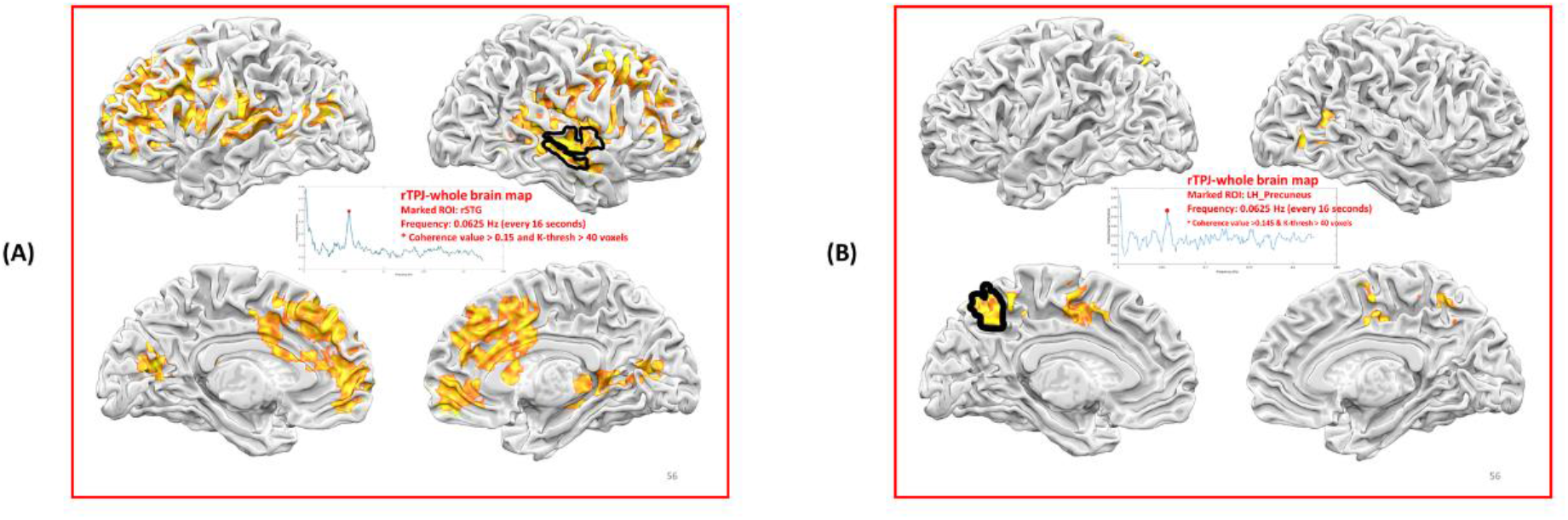
The rTPJ-whole brain frequency bin mapping. Strong coherence in each of the 224 frequency bins are shown. (A) Coherence is higher than 0.15 with a threshold over 40 voxels colored for the cooperation condition. The marked region is the right STG. Watch the video clip by clicking here: 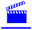. (B) Coherence is higher than 0.145 with a threshold over 40 voxels colored for the competition condition. The marked region is the left precuneus. Watch the video clip by clicking here: 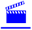.

**S5 Fig.**
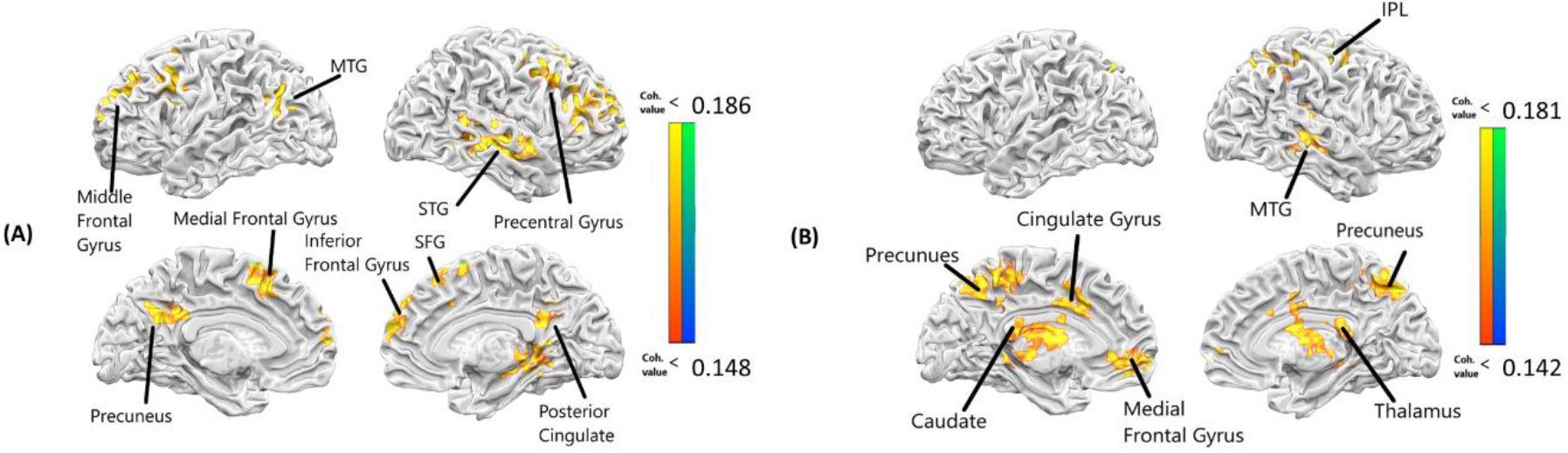
Brain regions showing significant coherence with the rSTG during the event of feedback time. (A) In the cooperation condition, under the cluster k-threshold of 40 voxels and the applied map threshold of 0.148, the ROIs with significant coherence with the rSTG in the cooperation condition include: the rSTG, the superior frontal gyrus (SFG), the right precentral gyrus, the right inferior frontal gyrus, the right posterior cingulate, the left medial frontal gyrus, the left precuneus, the left middle frontal gyrus, and the left middle temporal gyrus (MTG). (B) In the competition condition, under the cluster k-threshold of 40 voxels and the applied map threshold of 0.142, the coherent ROIs are: the bilateral precuneus, the right MTG, the right inferior parietal lobule (IPL), the right thalamus, the left caudate, the left cingulate gyrus, and the left medial frontal gyrus.

**S6 Table.** rSTG-Brain interpersonal coherence in cooperation and competition.

| **Cooperation condition with 448 beta volumes** |  |  |  |  |  |
| --- | --- | --- | --- | --- | --- |
| **Cluster** | **Anatomical region** | **x** | **y** | **z** | **coherence value** |
| **Cluster 1 (voxel: 61)** | L Medial Frontal Gyrus | -15 | 14 | 49 | 0.1858 |
| **Cluster 2 (voxel: 95)** | R Superior Frontal Gyrus | 9 | 59 | 28 | 0.1812 |
| **Cluster 3 (voxel: 150)** | R Superior Temporal Gyrus | 48 | -16 | -11 | 0.1810 |
| **Cluster 4 (voxel: 65)** | L Medial Frontal Gyrus | -12 | 59 | 13 | 0.1791 |
| **Cluster 5 (voxel: 45)** | L Precuneus | -6 | -49 | 31 | 0.1790 |
| **Cluster 6 (voxel: 61)** | R Superior Frontal Gyrus | 18 | 17 | 43 | 0.1765 |
| **Cluster 7 (voxel: 57)** | R Precentral Gyrus | 33 | 26 | 37 | 0.1749 |
| **Cluster 8 (voxel: 69)** | R Inferior Frontal Gyrus | 48 | 26 | 4 | 0.1728 |
| **Cluster 9 (voxel: 60)** | L Middle Frontal Gyrus | -51 | 11 | 37 | 0.1687 |
| **Cluster 10 (voxel: 50)** | L Middle Temporal Gyrus | -48 | -58 | 25 | 0.1681 |
| **Cluster 11 (voxel: 46)** | R Posterior Cingulate | 12 | -49 | 7 | 0.1679 |
| **Competition condition with 448 beta volumes** |  |  |  |  |  |
| **Cluster** | **Anatomical region** | **x** | **y** | **z** | **coherence value** |
| **Cluster 1 (voxel: 75)** | R Middle Temporal Gyrus | 54 | -31 | -5 | 0.1776 |
| **Cluster 2 (voxel: 154)** | R Precuneus | 12 | -64 | 49 | 0.1746 |
| **Cluster 3 (voxel: 117)** | L Caudate | -9 | 2 | 13 | 0.1717 |
| **Cluster 4 (voxel: 114)** | R Inferior Parietal Lobule | 33 | -28 | 43 | 0.1702 |
| **Cluster 5 (voxel: 60)** | R Thalamus | 3 | -34 | 8 | 0.1660 |
| **Cluster 6 (voxel: 57)** | L Precuneus | -15 | -46 | 37 | 0.1632 |
| **Cluster 7 (voxel: 41)** | L Medial Frontal Gyrus | -12 | 38 | -5 | 0.1599 |
| **Cluster 8 (voxel: 41)** | L Cingulate Gyrus | -3 | 11 | 31 | 0.1593 |

rSTG-Brain interpersonal coherence in the cooperation condition [Coherence value threshold > .148, cluster size > 40 voxels threshold] and in the competition during the feedback time with 448 beta volumes [Coherence value threshold > .142, cluster size > 40 voxels threshold].

**S7 Fig.**
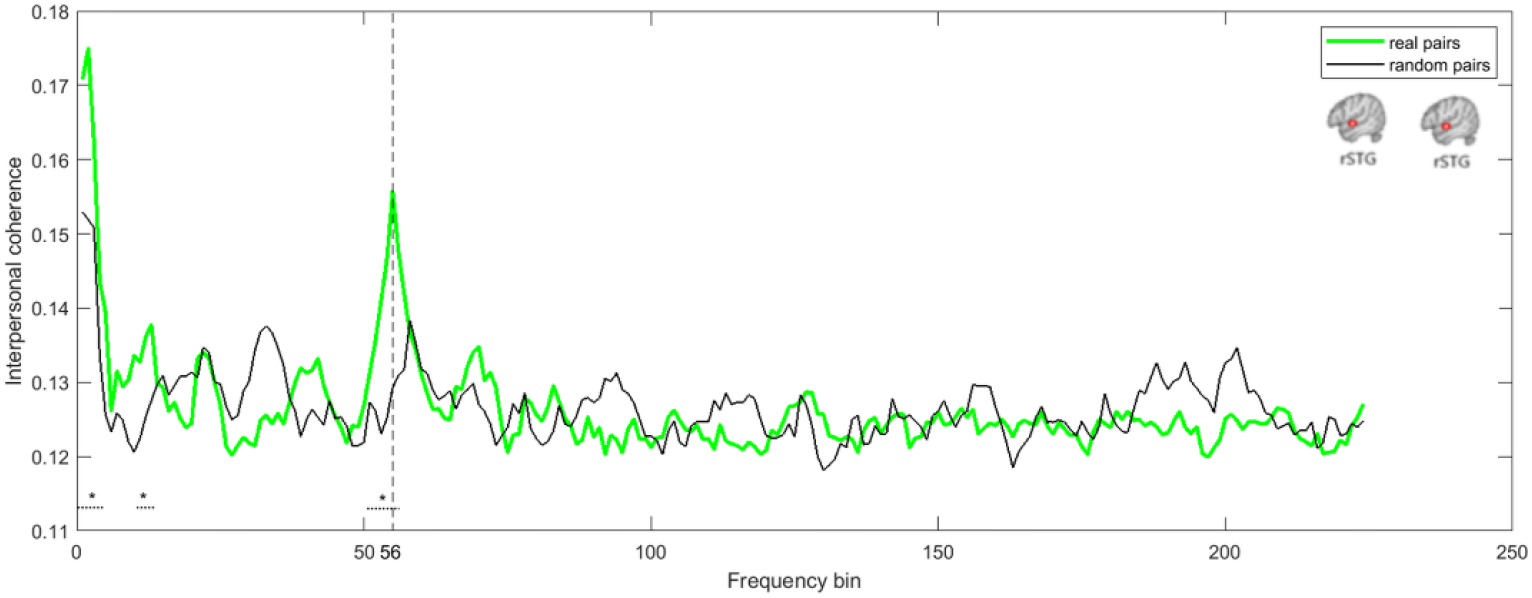
Interpersonal coherence spectra between the rSTG and the rSTG of real communicative pairs. The dominant (or peak) frequency, 0.0625 Hz (1/16 s), located at the 56^th^ frequency bin (along the x axis), revealed the 2^nd^ highest coherence (the first bins being around time zero). This bin corresponded to the average trial frequency (once every 17 s), as well as to the concatenated event frequency (beta series built by combining 8 betas, every 2 s/ 1 TR, after the trial feedback time); therefore, the heightened coherence between rSTG-rSTG suggest that the dyads reached certain degrees of synchronization (\**p* < 0.01 and three consecutive bins).

